# Single-Cell Proteomics of Human Peripheral Blood Mononuclear Cells Exceeding 600 Cells per Day

**DOI:** 10.64898/2026.02.01.703169

**Authors:** James M. Fulcher, Yumi Kwon, Pranav Dawar, Rashmi Kumar, Sarah M. Williams, Patricia Miller, Andrey Liyu, Liang Chen, Daniel J. Orton, Heather M. Olson, Fengchao Yu, Alexey I. Nesvizhskii, Julie Fortier, Ravi Vij, Reyka Jayasinghe, Li Ding, Ying Zhu, Ljiljana Paša-Tolić

**Affiliations:** Environmental Molecular Sciences Laboratory, Pacific Northwest National Laboratory, Richland, Washington, USA; Biological Sciences Division, Pacific Northwest National Laboratory, Richland, Washington, USA; Department of Pathology, University of Michigan, Ann Arbor, Michigan, USA; Gilbert S. Omenn Department of Computational Medicine and Bioinformatics, University of Michigan, Ann Arbor, Michigan, USA; Division of Oncology, Department of Medicine, Washington University School of Medicine, St. Louis, Missouri, USA; Department of Proteomic and Genomic Technologies, Genentech, Inc., South San Francisco, California, USA

**Keywords:** single-cell proteomics, PBMCs, real-time spectral library searching, TMTpro, nanoPOTS

## Abstract

Single-cell proteomic (scProteomic) measurements of peripheral blood mononuclear cells (PBMCs) are of considerable value in human health, given their involvement in the maintenance of healthy and diseased states. However, the high heterogeneity and relatively small size of immune cell types demand maximal throughput and sensitivity in proteomic measurements that have yet to be fully realized. Here, we describe an approach that addresses sensitivity and throughput through the implementation of Real-Time spectral Library Searching (RTLS), TMTpro 32-plex labelling, an updated nested-nanodroplet processing in One pot for Trace Samples (N2), and a dual-column liquid chromatography system. By prioritizing tandem mass spectrometry (MS2) features with high similarity to library spectra, RTLS enables greater identification depth and feature reproducibility than a standard shotgun MS2 approach in low-input and single-cell samples. The platform permitted 660 single PBMCs to be measured per day, with an average of 750 protein identifications per cell and 1,648 proteins in total, achieving the necessary throughput and depth to characterize immune cell populations. Application of this scProteomic method and a new cell typing informatics approach to 2,130 PBMCs enabled the identification of both major and low-frequency cell types (∼1-2%), as well as associated proteomic markers.

## Introduction

Peripheral blood mononuclear cells (PBMCs), a heterogeneous population comprising lymphocytes, monocytes, and dendritic cells, play central roles in immune surveillance, inflammation, and homeostasis in humans. Their composition and activation states are altered in numerous physiological and pathological conditions, including infection, autoimmunity, and cancer, making them a valuable resource for in-depth cellular characterization.^[1-2]^ To fully capture the cellular diversity and functional states of PBMCs, high-resolution and high-throughput single-cell approaches are essential.^[3]^

Single-cell RNA sequencing (scRNA-seq) has been transformative for dissecting the transcriptional landscape of PBMCs, enabling the identification of rare cell types, transient states, and lineage relationships.^[4-6]^ Its high throughput and scalability have made it a favored approach in immunology and clinical research. However, transcript abundance is often an imperfect proxy for protein abundance due to post-transcriptional/translational regulation, differences in mRNA and protein stability, and context-specific translation.^[7-8]^ This gap underscores the growing interest in single-cell proteomics as a complementary and more direct approach for understanding cellular responses, particularly in the context of immune-associated diseases.

Mass spectrometry-based single-cell proteomics (scProteomics) offers a promising approach for directly quantifying protein abundances across individual cells.^[9-10]^ Techniques such as nanoPOTS (nanodroplet Processing in One pot for Trace Samples)^[11]^ or nano-proteomic sample preparation (nPOP)^[12]^ have been developed to minimize sample losses and enabled the processing of single cells with picogram-level protein inputs. Such developments have been particularly necessary for PBMC cell types, as lymphocytes have a much smaller volume relative to cultured cells and the protein content is generally thought to scale with cell size^[13-14]^. As an example, the mean estimated volume of lymphocytes is ∼ 200 fL^[15-16]^ while HeLa cells are estimated to be ∼2,400 fL^[17]^ – representing an order of magnitude difference in the volume and therefore protein content.

Incorporation of TMTpro reagents into the aforementioned techniques (such as with the 16- and now 32/35-plex sets)^[18-19]^ enables multiplexed pooling of cellular material, thereby substantially increasing analysis throughput.^[20-22]^ In parallel, new developments have been made in liquid-chromatography (LC) efficiency through the use of dual-column systems,^[23-24]^ and recent intelligent data acquisition methods such as real time search (RTS) enhanced quant of single cell spectra (RETICLE) and prioritized Single-Cell ProtEomics (pSCoPE) have improved peptide identification efficiency and consistency.^[25-26]^ Yet, even with these developments, most studies typically only manage tens to ∼100 cells per day,^[27]^ with a single study reported to achieve ∼1,000 cells per day via LC-MS^[22]^. Therefore, scProteomic approaches still lag behind scRNAseq in terms of throughput and sensitivity, limiting its ability to make robust inferences in disease and cell typing.

Another recent development related to RTS on tandem ion-trap (IT) and Orbitrap instruments is real-time spectral library searching (RTLS).^[27]^ RTLS performs millisecond comparisons between experimental IT MS2 scans and library spectra to determine whether a given molecular feature should be selected for a higher-resolution MS2 acquisition. Multiplexed scProteomic methods often employ long ion injection times and time consuming high resolution MS2 acquisitions. Hence, the efficiency and overall depth of identified features could be improved by leveraging RTLS as a screen for high-quality features of interest.^[26]^ Furthermore, we sought to incorporate TMTpro 32-plex reagents into the nested-nanowell nanoPOTS approach,^[21]^ and combine this new platform with a dual-column system to significantly improve throughput and overall sensitivity for the analysis of PBMC cells. To this end, we demonstrate an updated N2 nanoPOTS platform for 32-plex TMTpro labeling of single PBMCs combined with a dual-column LC system. We further demonstrate that RTLS provides greater sensitivity and reproducibility in low-input and single-cells over standard MS2 approaches. Combining N2 nanoPOTS, a dual-column system, and RTLS enabled the analysis of 2,130 PBMCs (with an estimated median of 14.2 pg peptide input) over 77 hours of LC-MS acquisition time (∼660 cells per day) at a median identification depth of 752 proteins per cell.

## Results and Discussion

### Design and Development of the RTLS Approach

With a goal of performing high-throughput scProteomics on PBMCs, we envisioned an approach that incorporated: (1) N2 nanoPOTS for cellenONE-enabled, nanoscale preparation of TMTpro 32-plex labeled single-cells (**Figure 1a**), (2) a dual-column liquid chromatography system for more efficient sample trapping and peptide separation (**Figure 1b**), and (3) ion mobility (field asymmetric ion mobility spectrometry, FAIMS) mass spectrometry equipped with RTLS for improved selection of TMTpro-labeled peptide precursors during MS data acquisition (**Figure 1c**).

**Figure 1.**
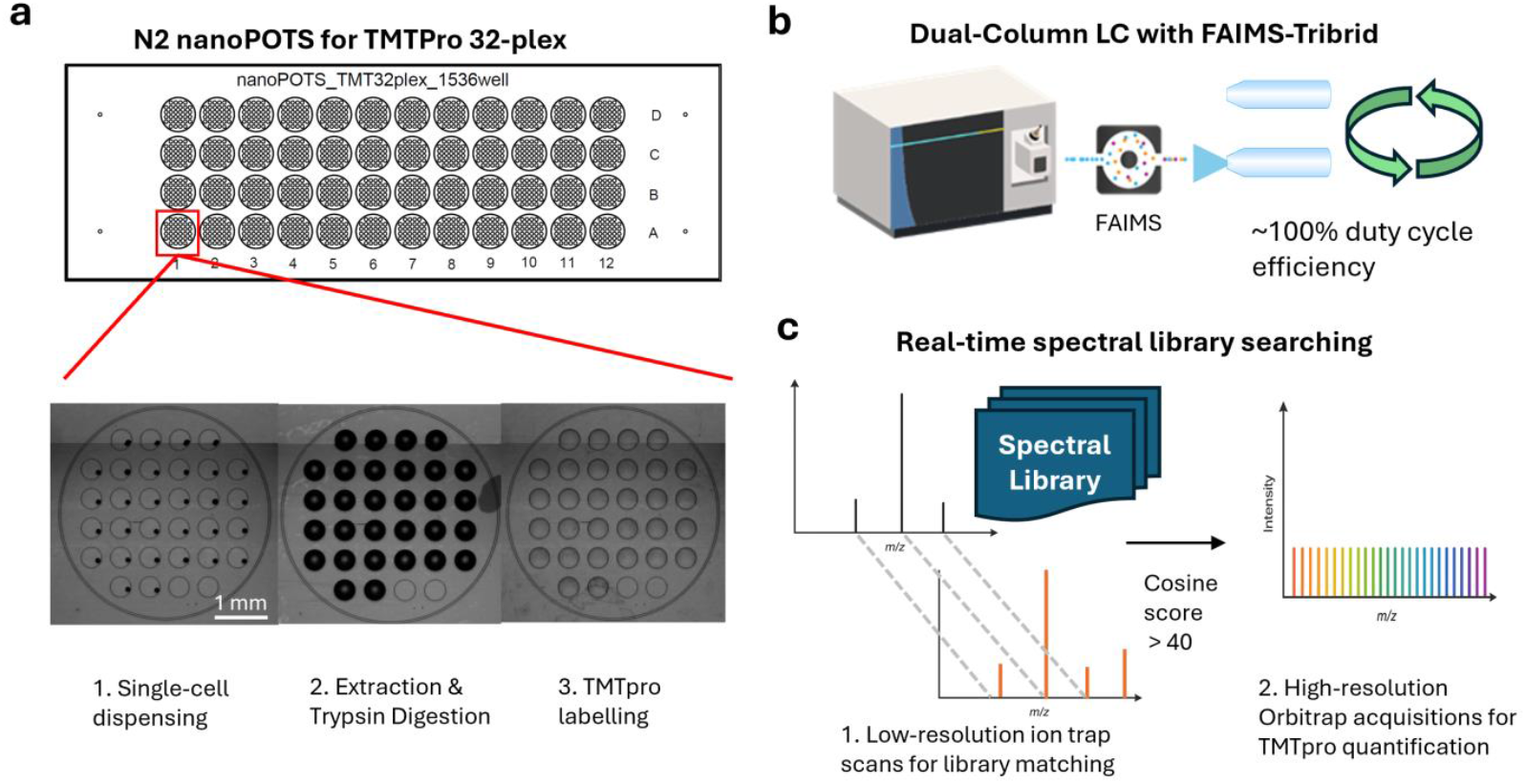
Overview of high-throughput scProteomics approach for analysis of PBMCs. (**a**) Design of the N2 nanoPOTS with 32 inner wells for compatibility with TMTpro 32-plex. Expanded image below shows example cellenONE-based processing steps on the chip, including single-cell dispensing, protein extraction and trypsin digestion, and TMTpro labelling. (**b**) Dual LC-column for efficient duty cycle during peptide separations with downstream MS analysis accomplished by FAIMS-equipped Orbitrap Eclipse Tribrid mass spectrometer. (**c**) RTLS approach showing library spectra matched to experimental spectral using low-resolution ion trap MS2 scans prior to high-resolution Orbitrap MS2 acquisitions for TMTpro 32-plex reporter ion quantification.

Towards this end, we implemented a new N2 device capable of incorporating TMTpro 32-plex reagents building upon our original 10-plex N2 nanoPOTS chip design.^[21]^ This new design incorporates 48 microwells nested with 32 inner nanowells thereby providing up to 1,536 positions for single-cell sample-processing per chip (**Figure 1a**). Additionally, we developed a dual-column LC system comprising two parallel LC setups to achieve nearly 100% mass spectrometry duty cycle. While the RTLS approach available on ThermoFisher Tribrid mass spectrometers has previously been demonstrated in bulk proteomic experiments,^[27]^ it has not been applied to low-input applications. Hence, we first evaluated the RTLS approach by comparing it to a “standard” single-cell MS2 approach using TMTpro-labeled PBMC peptide lysate (PBMCs isolated from a healthy human donor,^[28]^ labeled with channels 126 and 135ND) on our updated N2 nanoPOTS chips and dual-column LC system. To generate the spectral libraries for RTLS, the same TMTpro-labeled PBMC peptide lysate was fractionated and analyzed using both the ion trap and Orbitrap on a ThermoFisher Orbitrap Eclipse Tribrid mass spectrometer equipped with FAIMS operated at two compensation voltages (-45 and -65 V). This comprehensive library contained 184,253 spectra, 38,876 unique peptides, and 6,150 unique proteins.

We dispensed 10 ng of TMTpro-labeled PBMC peptide lysate onto a nanoPOTS chip and analyzed 10 replicates for the RTLS and “standard MS2” methods using equivalent MS instrument acquisition settings. We were encouraged by the finding that RTLS could provide 792 (9%) more unique peptide identifications across all datasets (**Figure S1a**), more peptides on average for each dataset (**Figure S1b**), and improved data completeness (**Figure S1b**) as compared with a standard MS2 approach. While 10 ng is at least an order of magnitude lower than typical bulk proteomic experiments, it is up to 250-fold higher than the amount of protein that might maximally be anticipated to be observed in a circulating lymphocyte (∼40 pg, assuming 20% w/v protein content for a cell of ∼200 fL volume).^[15-16, 29-30]^ Given the modest advantage of the RTLS in a 10 ng samples, and considering data sparsity and peptide observability are directly related to their abundance during MS analysis,^[31]^, we next applied RTLS to single PBMCs with the expectation that there is increasing benefit to RTLS with lower peptide input levels. As the RTLS method is restricted to selecting peptide features that are represented in the spectral library, we reasoned that RTLS should improve the overall number of peptide identifications in single-cell samples by ensuring features selected for the more time-consuming Orbitrap MS2 acquisitions have TMTpro modifications and identifiable fragment ions.

We sorted 30 PBMCs onto nanowells within six N2 nanoPOTS microwells (n = 3 for standard MS2 and RTLS), performed nanoscale sample preparation, and labeled with TMTpro. Normalization of the deuterated and non-deuterated TMTpro 16-plex was accomplished by including 250 pg of the 126/135ND-labeled PBMC lysate. The increase in unique and average peptide identifications was significantly higher for single-PBMC samples after applying RTLS in comparison to the 10 ng samples (**Figure 2**). RTLS yielded 685 (19%) more unique peptide identifications (**Figure 2a**), 526 (44%) more peptides on average for each single-cell (y-axis, **Figure 2b**), and significantly improved data completeness (x-axis, **Figure 2b**) as compared to the standard MS2 approach (26% standard MS2 vs 37% with RTLS). Taken together, these data demonstrate the advantages of RTLS in scProteomics. As a final optimization of the method, we applied the RTLS approach to several bridge inputs above and below the 250 pg used originally. The negative impacts of increasing amounts of TMT reference or bridge channel inputs on quantitative accuracy in scProteomics have been previously studied, and general recommendations suggest a maximum of 20-fold the expected single-cell input.^[32-34]^ We tested bridge channel inputs ranging from ∼1-fold up to 25-fold (assuming 40 pg per single-cell), measuring the unique peptides quantified from the bridge channel samples, single-cells, and “blank” (empty) TMTpro channels (**Figure 2c**). As expected the number of quantified peptides in bridge and single-cell channels showed diminishing returns between 250 and 1,000 pg inputs. Surprisingly, the blank channels (without peptides) also showed increasing number of peptide identifications with higher bridge-input levels. While this would be expected if the blank channels were isotopically adjacent to the bridge channels, the blank channels (134C, 135N, and 135CD) are isotopically non-adjacent. This suggests that while higher bridge inputs provide more ions for precursor detectability and MS2 fragmentation, low *m/z* peptide fragment ions can cause interference even with high-resolution Orbitrap acquisitions and 10 ppm mass accuracy during reporter ion quantification (a result similar to previous isobaric, low-input samples analyzed with low-resolution linear ion trap).^[35]^ Our results suggest that bridge channel inputs should remain below 1,000 pg (< 25-fold), consistent with prior work.^[32-34]^

**Figure 2.**
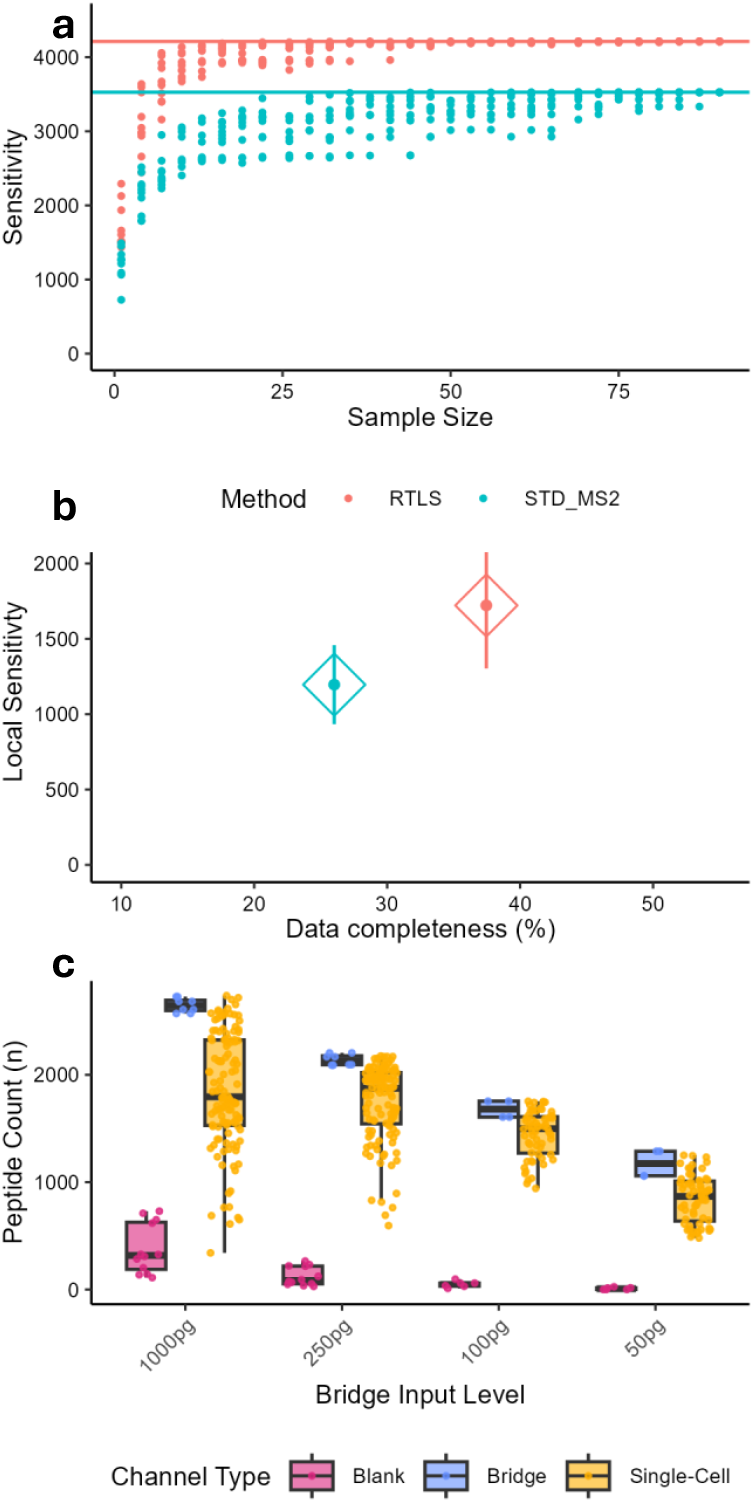
Comparison between RTLS and standard MS2 method using single PBMCs (n = 90 cells/ n = 3 TMT batches). (**a**) Cumulative sensitivity curves showing peptides identified as datasets are sampled. Solid line indiciates total sensitivty (total unique peptides) (**b**) Mean local sensitivity (average number of peptides per dataset) compared to overall data completeness. Lines represent +/-1 standard deviation. (**c**) Boxplots of different bridge input levels with single PBMCs; 1000 pg (n = 120), 250 pg (n = 120), 100 pg (n = 60), and 50 pg (n = 60). Lines indicate median and boxes specify interquartile range (IQR)

### Application of the RTLS Approach to Thousands of PBMCs

With the optimizations of the RTLS approach in place, we sorted and prepared single PBMCs with TMTpro 32-plex reagents on the new N2 nanoPOTS chips followed by the addition of 300 pg of bridge sample divided equally in channels 126 and 135ND (150 pg each). Acquisition of 71 TMTpro batches representing 2,130 cells was achieved using a dual-column system coupled to an Orbitrap Eclipse Tribrid mass spectrometer, with nearly 100% MS utilization. Collection of this data corresponded to a throughput of approximately 660 cells per day. Database searching with the FragPipe proteomics pipeline^[36-43]^ and TMTpro 32-plex quantification using TMT-Integrator^[44]^ afforded a median of 2,164 peptide and 752 protein identifications per single-cell (**Figure 3a, 3b**, and **Table S1**). As expected, blank channels showed markedly lower numbers of quantified peptides and proteins than single-cell channels, and bridge channels generally provided deeper coverage than single-cell channels. Summing of the reporter ion intensities for each sample and calculating the medians for the blank, bridge, and single-cell channels demonstrates a 10.5-fold difference between the bridge and single-cell samples, and a 380-fold difference between the single-cell samples and blank channels (**Figure 3c**). As each bridge channel contained 150 pg of peptide input, we back-calculated the median single-cell:bridge channel ratio to estimate the amount of peptide input detected for each cell (**Figure 3d**). Our data suggests that the median PBMC provided approximately 14.2 pg of peptide input, demonstrating the approach’s sensitivity to smaller cells.

**Figure 3.**
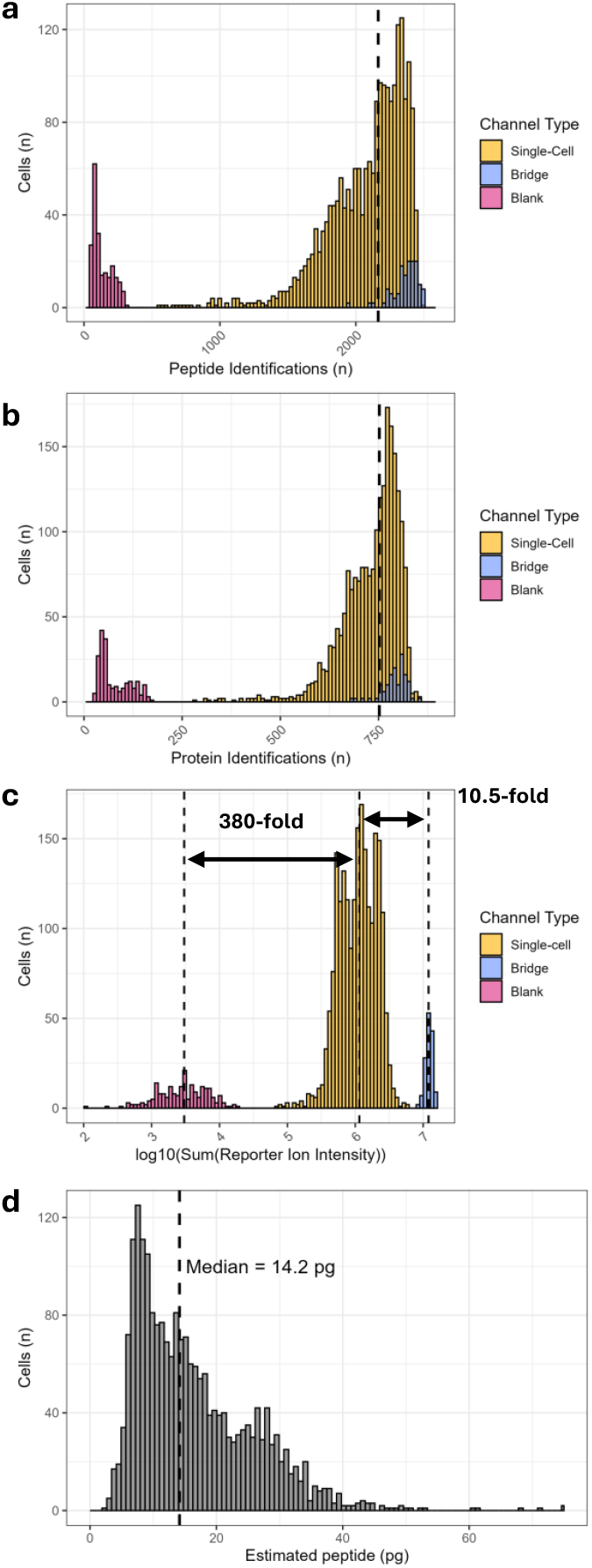
Depth of identifications and TMTpro reporter ion signal in single PBMCs (n = 2130), blank (n = 213), and bridge (n = 142) channels. (**a**) Histograms of peptide identifications, (**b**) protein identifications, and (**c**) log_10_ mean reporter ion intensities. Dashed lines indicate medians of designated distributions. (**d**) Histogram of estimated peptide input (pg) from each single-cell.

We next implemented an informatics pipeline that provided greater insights into the cell types detectable in our single-cell datasets. First, to minimize confounding effects in downstream analysis, we performed median normalization and correction (using ComBat) of known batch effects that are a result of sample preparation and LC-MS analysis (**Figure S2**).^[45]^ Imputation (for analyses requiring complete data) yielded 1,648 unique, quantified proteins **Table S2**). Next we leveraged Seurat’s multimodal reference mapping capabilities to use scRNAseq data as a guide for cell typing of scProteomics data.^[46]^ A large subset (20,000) of unstimulated PBMCs was randomly sampled from a recent scRNAseq dataset published by Oelen et al.^[5]^ and used to transfer coarse, high-level (i.e., CD4 T-cell, monocyte, NK cell, etc…) labels to our scProteomics data. In the first pass through this pipeline, we noted both CD4+ T-cells (referred to as CD4T_1 and CD4T_1_2) and monocytes (referred to as monocyte_0 and monocyte_0_2) were present in two distinct clusters, respectively, in the UMAP embedding of the scProteomics data (**Figure S3**). We believe this likely reflects a recently identified phenomenon in scProteomics in which single cells can lose protein content to leakage from unintentional permeabilization during cell sorting.^[22]^ This inference relies on several key observations, specifically: (1) these bifurcated clusters show distinct estimated peptide distributions derived directly from non-median normalized TMTpro reporter ion intensities (**Figure S4A**), (2) total protein identifications per single-cell are higher in one cluster than the other (**Figure S4B**), (3) the distributions of maximum Seurat prediction scores show distinct differences between clusters (**Figure S4C**), and (4) gene ontological terms representing proteins decreased in abundance (higher peptide input cluster vs lower peptide input cluster) show shared enrichment of terms related to cytosolic proteins (**Figure S5**).Collectively, this data indicates that one cluster of cells for both CD4 T-cells and monocytes had lower protein content, and this protein content was biased towards cytosolic proteins which we interpret as relating to cell permeabilization prior to isolation. Therefore, to ensure the integrity of the downstream cell typing and quantitative comparisons, cell clusters (specifically CD4T_1_2 and monocyte_0_2) with the lower protein content were removed providing 1,275 single-cells for further analysis.

As our pipeline implements Seurat multimodal reference mapping, one could in principle assign cell identities directly from the cell prediction scores. However, we found that even with these 1,275 higher quality cells, 249 still fell below a Seurat prediction score cutoff of 0.56 (**Figure S6**). Note that we defined this cutoff as the mean prediction score minus 1 standard deviation, following prior work on construction of single-cell reference atlases for human and mouse lung.^[47]^ With multimodal reference mapping, a caveat in mapping unrelated modalities and datasets is the reliance on overlapping features and the assumption that these features will exhibit correlated variation between modalities. In our data, 188 features overlapped between the 600 highest variable proteins in our scProteomic dataset and 2,000 highest varying genes in the scRNAseq atlas (**Figure S7**). Thus, a large fraction of the proteome-level heterogeneity is “invisible” to the mapping procedure as the reference

Though we were encouraged by the presence of overlapping immune-relevant features with high variance, including TLR2 (toll-like receptor 2), GZMB, GZMM, and CDA (**Figure S7**), we sought a hybrid strategy that would also leverage the many highly variable proteins that are unique to the scProteomic data. Therefore, we computed the shared nearest neighbor graph from the PCA of the 600 highest varying proteins in our dataset followed by Louvain clustering to identify clusters based purely on the scProteomic data (**Figure S8a**) and then identified which Seurat predicted cell types (**Figure S8b**) were statistically enriched for a given cluster using the hypergeometric test (**Figure S8c**). The test uses only high-confidence cells (prediction score ≥ 0.56) but the resulting cluster-level identity is propagated to all cells in that cluster (see **Supplementary Methods and Experimental Procedures** for more detail). Reanalysis of these cells using our cell-typing pipeline provided eight major clusters from which six different cell types could be identified (**Figure 4a**). Of the 1,275 high-quality cells, 1,251 were assigned to one of six major cell types while 24 cells could not be confidently assigned a cell type (referred to as “unknown”). Assigned cell types included monocytes (n = 456), CD4+ T-cells (“CD4T”, n = 308), CD8+ T-cells (“CD8T”, n =181), natural killer cells (“NK”, n = 150), B-cells (“B”, n = 102), and dendritic cells (“DC”, n = 54). Notably, monocytes could be represented as two clusters.

**Figure 4.**
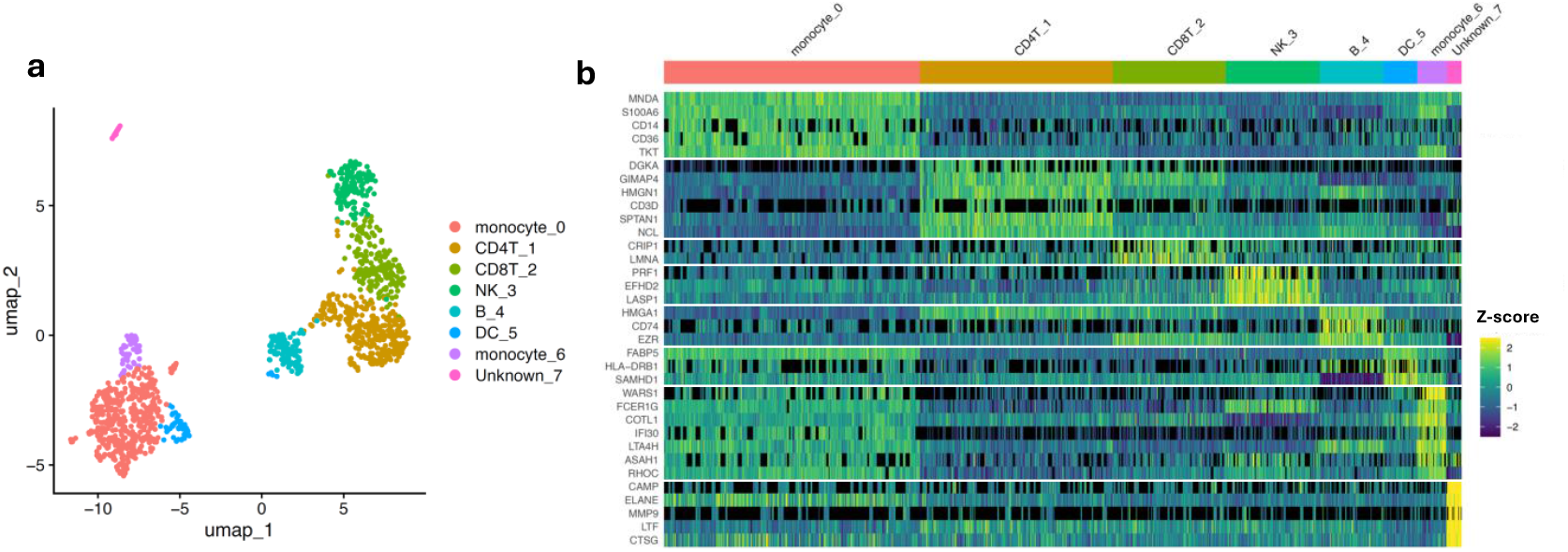
Cell types and marker proteins identified from our hybrid annotation approach (**a**) UMAP generated from PCA of the 600 highest varying proteins from the scProteomic data. Final cell types assigned are colored as indicated. “DC” = dendritic cell, “NK” = natural killer cell, and “B” = B-cell. (**b**) Heatmap of the most statistically significant proteins enriched for each cell type, as well a selected known markers. Black coloring indicates missing values while viridis color indicates relative protein abundance (Z-score). Statistical significance was determined through a two-sided limma/Empirical Bayes test with Bonferroni correction of p-values for multiple hypothesis testing.

We performed differential abundance analysis between these cluster-aware cell types to identify enriched proteins (**Figure 4b** and **Table S3**). Encouragingly, many statistically significant proteins reflected known markers. Myeloid cell nuclear differentiation antigen (MNDA) is a nuclear protein expressed in myeloid-lineage cells such as monocytes.^[48]^ GTPases of the immunity associated proteins (GIMAPs, such as GIMAP4) and diacylglycerol kinases (such as DGKA) are associated with T-cell differentiation, development, and function at maturity.^[49-51]^ CD74, the receptor for macrophage inhibitory factor, is an established B-cell marker and regulator of transcription in B-cells.^[52]^ HLA-DRB1 is a classical major histocompatibility complex class II (MHC-II) protein enriched on dendritic cells, aligned with their functions as antigen-processing and presenting cells.^[53]^ Other canonical immune markers such as CD14 (monocytes), CD36 (monocytes), and CD3D (T-cells) were also found enriched in their respective cell types (**Figure 4b** and **Table S3**). Interestingly, monocytes were found to occupy two distinct clusters. The differential abundance analysis suggests the smaller cluster (monocyte_6) represents intermediate (CD14+/CD16+) or non-classical (CD14dimCD16+) monocyte subtypes. This is supported by the greater abundance of several inflammation-related proteins (**Figure 4b**); including WARS1, FCER1G (gamma chain of high affinity IgE receptor), LTA4H (Leukotriene-A4 hydrolase, a proinflammatory enzyme),^[54]^ ASAH1 (acid ceramidase, recently found to regulate innate immunity),^[55]^ and IFI30 (Interferon Gamma-Inducible Lysosomal Thiol Reductase) - all of which show similar patterns of enrichment for CD16+ monocytes in scRNAseq datasets (**Figure S9**).^[56]^ Additionally, cell-motility related proteins indicative of non-classical monocytes (specifically COTL1 and RHOC) were found to be enriched in this cell type (**Figure 4b**).^[57]^

Finally, the “unknown” cell type demonstrates the advantage of our hybrid annotation approach that leverages both the underlying scProteomic features and scRNAseq mapping results (**Figure 4a** and **4b**). If we had relied solely on the Seurat-predicted annotations, this small cluster of 24 cells would have either been left out of further analysis or incorrectly grouped with other cell types (**Figure S8b**). Instead, our clustering-informed approach enabled us to identify several highly enriched proteins specific to this group of cells, all of which (CAMP, ELANE, CTSG, MMP9, and LTF) are hallmark proteins associated with granulocytes/neutrophils, a cell type whose presence is likely a result of unintentional contamination during PBMC isolation.^[58-61]^ These unique cells have notable relevance to many diseases and have only recently been analyzed through MS-based scProteomic approaches.^[62]^ Taking these results together, this work demonstrates the exceptional throughput and reproducibility of the TMT 32-plex N2 chip and dual-column workflow when integrated with RTLS. Through the analysis of a large population of PBMCs, we also validate canonical immune markers and identify new potential protein-specific markers in PBMC cells that may be unresolvable with other -omics approaches.

## Conclusion

In this study, we demonstrate that integrating Real-Time spectral Library Searching (RTLS) with TMTpro 32-plex labeling, N2 nanoPOTS sample preparation, and a dual-column LC-MS platform substantially advances the throughput, sensitivity, and reproducibility of scProteomic measurements. RTLS consistently improved peptide identification depth and data completeness in both low-input and single-cell PBMC samples, highlighting the value of real-time targeting of reproducible precursor features for Orbitrap MS2 acquisition. The combined workflow enabled the analysis of 2,130 PBMCs in 77 hours - achieving a throughput of 660 cells per day while maintaining a median of ∼750 proteins identified per cell.

By further coupling this experimental platform with a hybrid informatics cell-typing strategy that leverages both proteome-driven clustering and multimodal reference mapping to scRNAseq atlases, we resolved six major circulating immune cell types and detected low-frequency dendritic cells, a monocyte subtype, and granulocytes with corresponding differentially abundant protein signatures. These results illustrate the power of isobaric scProteomics to capture cell-type-specific proteomic signatures at high throughput while also quantifying markers consistent with established immune biology. Through the approach presented here, we anticipate that scaling the number of cells ten-fold with similar scaling of biological replicates will enable the identification of new protein therapeutic targets in the context of immune-mediated diseases such as myelomas, lymphomas, and leukemias.

Overall, this work establishes an accessible framework for large-scale scProteomic profiling of heterogeneous immune samples. The improvements in sensitivity, consistency, and throughput enabled by RTLS and TMTpro 32-plex nanoPOTS demonstrate scProteomics as a robust and increasingly mature counterpart to scRNAseq approaches, with potential for applications in immunology, disease profiling, and biomarker discovery.

## Supporting Information

The authors have cited additional references within the Supporting Information.^[5, 11, 28, 56]^

## Acknowledgements

We thank Dr. Matthew Monroe for his assistance in depositing the raw proteomic data onto MassIVE. We also thank William Barshop and Rafael Melani (ThermoFisher Scientific) for assistance in implementing the RTLS approach. This work was supported by the National Institutes of Health Grant R01CA272377. A portion of the research was performed using the Environmental Molecular Sciences Laboratory (grid.436923.9), a DOE Office of Science User Facility sponsored by the Biological and Environmental Research program under Contract No. DE-AC05-76RL01830.

## Data Availability Statement

The mass spectrometry raw data and database search results are accessible through the ProteomeXchange Consortium and MassIVE data repository with dataset identifiers MSV000100682 (PXD073879, PBMC bridge channel fractionated data), MSV000100683 (PXD073880, 10 ng and single PBMC optimization samples) and MSV000100684 (PXD073881, scProteomics of 2,130 PBMCs). Associated code for the data analysis, result tables, single-cell images, and metadata used for figure generation can be found at the GitHub repository cited here.^[63]^

## Entry for the Table of Contents

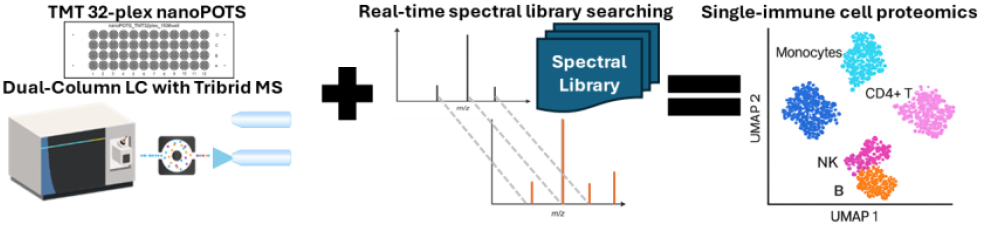

A high-throughput single-cell proteomics workflow integrating Real-Time spectral Library Searching (RTLS), nested-nanodroplet processing (N2 nanoPOTS), and TMTpro 32-plex quantification enables deep profiling of over 2,000 peripheral blood mononuclear cells (PBMCs), revealing major and rare immune cell types with proteome-level resolution.

