## Supplementary Information for "Single-Cell Proteomics of Human Peripheral Blood Mononuclear Cells Exceeding 600 Cells per Day"

### Table of Contents

#### Methods and Experimental Procedures

Materials ……......….......…………………………………………………...………………3
Fabrication of nanoPOTS chips……………………..…....…...…………...…..……………...3
Bulk processing of PBMC samples………......…………………………………………………4
TMTpro labeling of PBMC bridge sample…......…………………………………………………5
Micro-fractionation of TMTpro labeled PBMC bridge sample………......……………………5
Single-cell isolation on cellenONE ………………………………............…......………..…..6
Database Searching……………………………………………………………………......11
Spectral Library Generation for RTLS…………………………………………………...…12
Data Importing, Normalization, and Batch Correction Steps…………………………...…...13
Seurat Multimodal Reference Atlas Construction…………………………………………......14
Seurat Multimodal Annotation Transfer……………………………………………..…...15
Proteomics-driven dimensionality reduction and clustering………………...………15
Differential Abundance and Gene Enrichment Analysis…………………....…………….16

#### Tables

Table S1. Median-normalized, FragPipe output for all single PBMCs……….…....…17

Table S2. Single PBMC median-normalized, batch-corrected, and imputed matrix…17

Table S3. Differential abundance results across assigned cell types…………….....…17

Table S4. Table listing the datasets and MS methods performed in this study………18

Table S5. Analytical Metadata for all single PBMCs ………………………….……..……18

#### Figures

Figure S1. Comparison of RTLS and standard MS2 method using 10 ng input …….20

Figure S2. Eigencor plots showing the correlation of batch effects with PCs………..21

Figure S3. UMAP of identified cell types from initial cell typing analysis…..…………22

Figure S4. Histograms comparing estimated peptide input, identified proteins, and Seurat prediction scores for CD4+ T-cell and monocyte clusters. ………………...…...23

Figure S5. Gene enrichment analysis of CD4+ T-cell and monocyte clusters……….24

Figure S6. Histogram of Seurat prediction scores for 1,275 PBMCs……...………….25

Figure S7. Variable feature plot of 1,648 proteins measured in 1,275 PBMCs……...26

Figure S8. Louvain clustering, Seurat predicted, and final annotated cell types…….27

Figure S9. scRNAseq expression data for CD16+ monocyte enriched proteins…….28

**References…………………………………………..…………………………………..….28**

### Methods and Experimental Procedures

#### Materials

Deionized water (18.2 MΩ) was purified using a Barnstead Nanopure Infinity system (Los Angeles, CA, USA). Calcium chloride, N-dodecyl-β-D-maltoside (DDM) and formic acid (FA) were obtained from Sigma (St. Louis, MO, USA). Trypsin platinum and Lys-C were purchased from Promega (Madison, WI, USA). TMTpro 32-plex reagents (96-well plate format), dithiothreitol (DTT, No-Weigh format), 0.5 M Bicine solution, anhydrous DMSO, 50% aqueous hydroxylamine, LC-MS grade acetonitrile (ACN) with 0.1% FA, and LC-MS grade water with 0.1% FA were acquired from Thermo Fisher Scientific (Waltham, MA, USA). Polyimide Kapton tape was acquired from 3M (Maplewood, MN, USA).

#### Fabrication of nanoPOTS chips

The N2 nanoPOTS chips were fabricated using standard photolithography, wet etching, and silanization as described previously [11]. Each chip contained 48 (4 x12) nested nanowells with a diameter of 4 mm encircling 32 inner wells with diameters of 0.4 mm. Distances between inner wells are 0.55 mm while nested interwell distances are 4.5 mm. Chip fabrication utilized a 25 mm x 75 mm glass slide pre-coated with chromium and photoresist (Telic Company, Valencia, USA). After photoresist exposure, development, and chromium etching (Transene Company, Inc.), the exposed region was etched to a depth of ~5 μm with buffered hydrofluoric acid. The freshly etched slide was dried by heating it at 120 °C for 2 h and then treated with oxygen plasma for 3 min (AP-300, Nordson March, Concord, USA). 2% (v/v) heptadecafluoro-1,1,2,2-tetrahydrodecyl-dimethylchlorosilane (PFDS, Gelest, Germany) in 2,2,4-

trimethylpentane was applied onto the chip surface and incubated for 30 min to allow for silanization. The remaining chromium covering the wells was removed with chromium etchant, leaving elevated hydrophilic nanowells surrounded by a hydrophobic surface. Polyetheretherketone (PEEK) chip covers were machined as chip covers to provide sealing and protection of the droplets from evaporation during sample processing. Chips were wrapped in aluminum foil and parafilm for long-term storage and intermediate steps during sample preparation.

**PBMC sample isolation and bulk processing**

Blood samples taken from healthy donors were processed following previous protocol[28] and stored under liquid nitrogen until use. Two 1 mL aliquots of healthy donor PBMC samples (3.6 million cells per mL) were washed as described above to produce a cell pellet. To this pellet, 0.2 mL of 8 M Urea in 50 mM triethylammonium bicarbonate (TEAB) was added. Sample was briefly vortexed, pipetted up/down to lyse cells, and then incubated at 37°C for 1 hr with shaking at 850 RPM in a ThermoMixer. A bicinchoninic acid protein assay (BCA) was then performed on the lysate and determined to be 0.448 μg/μL (89.6 μg total). Sample was diluted 8-fold with 1.14 mM $CaCl_2$ in 50 mM TEAB (1.6 mL total volume). Lys-C was added at a ratio of 1:50 (Lys-C:Sample Protein) and incubated for 2 hr at 25°C with 850 RPM shaking. Next, trypsin was added at a ratio of 1:50 followed by overnight (14 hr) incubation at 25°C with shaking at 850 x RPM.

For solid-phase extraction cleanup, a C18 MacroSpin column (Nest Group) was prepared according to manufacturers' instructions. The digested sample was acidified by adding trifluoroacetic acid (TFA) to a final concentration of 0.1% TFA, and then loaded onto the MacroSpin column in 400 µL increments using 100 x G centrifugation for 1 min. The sample was then washed with 4 x 400 µL volumes of 5% ACN with 0.1% TFA and eluted with 400 µL of 80% ACN, 0.1% TFA. Sample was then concentrated to 30 µL using a SpeedVac before a final BCA assay was performed to determine peptide concentration (53 µg per 30 µL). Peptide was then dried to completion before further processing.

**TMTpro labeling of PBMC bridge sample**

To achieve TMTpro labeling of bulk PBMC peptide sample, the sample was reconstituted with 53 µL 50 mM bicine pH 8.5. 10 µg of peptide (10 µL) was then transferred to two different Eppendorf tubes before addition of 5 µL of TMTpro reagent (Channel 126 for one tube and channel 135ND for the other) in anhydrous DMSO. Incubation at room temperature was performed for 1 hr prior to quenching of the reaction through addition of 1 µL of 5% hydroxylamine followed by another 1 hr incubation. The two TMTpro labeled bridge channels were then combined into a single Eppendorf tube and diluted to a concentration of 0.2 µg/ µL through addition of 70 µL of 50 mM bicine.

**Micro-fractionation of TMTpro labeled PBMC bridge sample**

TMTpro-labeled peptide sample (0.2 µg/ µL) were separated by high resolution reverse phase cLC using a nanoACQUITY UPLC® system (Waters Corporation, Milford, MA) equipped with an autosampler. Capillary columns, 200 µm i.d. × 65 cm long, were packed with 300 Å Jupiter C18 bonded particles (Phenomenex, Torrence, CA). Separations were performed at a flow rate of 2.2 µL/min on binary pump systems, using

10 mM ammonium formate (pH 7.5) as mobile phase A and 100% acetonitrile as mobile phase B. 20 µL of TMT labeled peptide mixtures (0.5 µg/µL) were loaded onto the column and separated using a binary gradient of 1% B for 35 min, 1-10% B in 2 min, 10-15% B in 5 min, and 15-25% B in 35 min, 25-32% B in 25 min, 32-40% B in 13 min, 40-90% B in 43 min, held at 90% B for 2 min, 90-50% B in 2 min, 50%-95% B in 2 min, held at 95% B for 2 min and 95-1% in 4 min. The capillary column eluent was automatically deposited every minute into 12 x 32 mm polypropylene vials (Waters Corporation, Milford, MA) starting at minute 65 and ending at minute 170, over the course of the 180 min LC run. Fractions were concatenated by collecting fractions from vial 1 to vial 6 and then returning to vial 1 back to vial 6 (etc.). Prior to peptide fraction collection, 100 µL of water was added to each vial to aid the addition of small droplets into the vial.  Each vial was completely dried in a vacuum concentrator (Labconco, Kansas City, MO) and reconstituted in 15 µL 2% acetonitrile, 0.1% formic acid.

**Single-cell isolation on cellenONE**

Healthy donor PBMC samples (3.6 million cells per mL) stored in 1 mL cryopreservation media (12.5% human serum albumin, 10% DMSO) were thawed in a 37°C water bath until a limited amount of ice remained. Cells in cryopreservation media were then added to 9 mL of pre-warmed PBS (37 °C) in a 15 mL falcon tube, disinfected with 70% EtOH in BSL-2 cabinet, and transferred to a centrifuge for centrifugation at 250 x G for 10 minutes (rt). The falcon tube containing cell pellet was transferred to the BSL-2 hood prior to decantation and gentle resuspension of the pellet in 10 mL of pre-warmed PBS. The above washing process was repeated twice before cells were resuspended in 1.5 mL of PBS to a final concentration estimated at 2.4 million cells per mL.

PBMCs were then transferred to a 400 µL Eppendorf and placed in a cellenONE instrument equipped with a glass piezo capillary (P-20-CS) for dispensing and aspiration. Sorting parameters included a pulse length of 50 µs, a nozzle voltage of 79 V, a frequency of 500 Hz, a LED delay of 200 µs, and a LED pulse of 3 µs. The slide stage holding the N2 nanoPOTS chips was operated at dew-point control mode to reduce droplet evaporation. Cells were isolated based on their size, circularity, and elongation in order to exclude apoptotic cells, doublets, or cell debris. Specific light transmission isolation parameters included cell diameters between 15 and 22 µm, minimum threshold intensity of 20, circularity maximum of 3, and elongation maximum of 2.4. Single PBMCs were sorted into 30 of the inner wells of the 32-plex N2 nanoPOTS chips, apart from 2 wells which were left empty for addition of bridge sample. After sorting, N2 chips were wrapped in aluminum foil and parafilm prior to storage at -80 °C.

**N2 nanoPOTS Single-Cell Sample Preparation**

On the day of sample preparation, the N2 chips were removed from the freezer and allowed to thaw at room temperature before being moved onto ice. N2 chips were placed back in the cellenONE with the cooler set to -1.5 °C below dewpoint to reduce droplet evaporation. For each well containing single-cells, 10 nL of TMT-compatible digestion buffer (25 mM bicine, 25 mM NaCl, 10 mM $CaCl_2$, 0.05% DDM with 0.005 ng/nL Lys-C and 0.01 ng/nL trypsin) was added. Chips were dispensed one at a time and immediately covered in parafilm and aluminum foil before being placed in a high-humidity container in a 37 °C incubator for overnight digestion. The following day, the chips were removed from the 37 °C incubator and placed back in the cellenONE with the cooler set to room temperature (22 °C). A 96-well plate containing TMTpro 32-plex reagents was brought to room temperature as well. Anhydrous DMSO was placed in the cellenWASH position of the cellenONE and each TMTpro channel (50 µg in 5 µL stabilized DMSO solution) was diluted to 30 µL (1.67 ng/nL) with anhydrous DMSO immediately prior to dispensing. 25 µL of each channel was transferred to a 384 well plate inside the cellenONE for dispensing, and 5 µL of DMSO was drawn up into the PDC followed by 5 µL of TMTpro. Field files were adjusted to the appropriate position and 1 nL (1.67 ng) of the TMTpro solution was dispensed onto the well. After dispensing, PDC was flushed before repeating for each TMTpro channel. Once all channels were dispensed, the humidity in the cellenONE was increased to 60% and the deck temperature was set to 18 °C before overnight incubation. The TMTpro labeling reaction was then quenched through addition of 1 nL of 1 % hydroxylamine in LC-MS grade water with 2 mM dithiothreitol followed by a 30 min incubation. Finally, 5 nL of 5% formic acid was used to acidify the sample followed by evaporation of the remaining liquid and storage at -20 °C. TMTpro labeled bridge sample (0.02 ng/nL) was added at amounts as indicated in the main figures to each of the 32-plex wells prior to LC-MS analysis.

**Liquid Chromatography Methodology**

A homebuilt dual-column nanoPOTS autosampler was employed to automatically perform sample collection, cleanup, and liquid chromatography (LC) separation. The dual-column system consisted of two parallel LC setups, each comprising an in-house packed solid-phase extraction column (100 µm i.d., 4 cm, packed with 5 µm, C18 packing material (300 Å pore size; Phenomenex), and an in-house packed LC column

(50 μm i.d., 25 cm-long, packed with 1.7 μm, C18 packing material (BEH 130 Å C18 material, Waters) heated at 50 °C with a column heater (Analytical Sales and Services Inc). The dual-column configuration was controlled by proprietary software to allow near-continuous operation by alternating between columns, utilizing one column for performing peptide separation while the other simultaneously equilibrated and loaded sample achieving a ~100% mass spectrometer duty cycle. LC pumps included two UltiMate 3000 RSLC nano Systems (Thermo Fisher Scientific).

Dried peptide samples on chips were dissolved with buffer A (0.1% FA in water), and then injected into the solid-phase extraction column for 5 min with a loading buffer containing 2% acetonitrile. After washing, samples were eluted and separated at 100 nl/min using gradient of buffer B (0.1% formic acid in acetonitrile). A 60-min separation from 8% to 22% buffer B was performed followed by a 15-min linear gradient from 22% to 35% mobile phase B. To introduce high voltage to the column, a custom-designed clamp was used. The clamp was modified to include two emitter tips, enabling voltage application to both LC columns in the dual-column system during alternating operation.

**Mass Spectrometry Methodology**

For mass spectrometry analysis, a ThermoFisher Orbitrap Eclipse Tribrid mass spectrometer with field asymmetric ion mobility spectrometry (FAIMS) was utilized. Depending on the exact set of data, different methods were employed. Common to all methods, however, were a spray voltage of 2400 V, ion transfer tube temperature of 200 ºC, FAIMS carrier gas flow of 4 L/min, and data collection time of 60 minutes. All other methodological details with dataset-specificity are described below (dataset and method metadata are described in **Table S4**):

<u>Method A (Standard MS2 Method):</u> In this method ionized peptides were fractionated by the FAIMS interface using internal compensation voltage stepping (-45 and -65 V) with a cycle time of 2.5 s for -45 V and 1.5 s for -65 V. Fractionated ions within a mass range 400-1400 m/z were acquired at 120,000 resolution with a max injection time of 50 ms, AGC target of 1E6, and RF lens of 30%. MIPS mode was enabled and precursor fit was set to 70% with a fit window of 1.2 m/z. Precursor ions were isolated with a window of 1.2 m/z, fragmented with 35% normalized HCD collision energy and analyzed in the Orbitrap with an AGC target of 250,000, a maximum injection time of 500 ms, and tandem MS2 data acquired starting at 110 m/z.

<u>Method B (RTLS Method):</u> All the above settings of Method A were used in this method, apart from additional parameters specific to the RTLS approach. In this RTLS approach, initial tandem MS2 scans are recorded using the ion trap and searched against previously made spectral libraries (one for each FAIMS CV). Precursor ions for spectral library searching were isolated with a window of 1.2 m/z, fragmented with 35% normalized HCD collision energy, and analyzed in the ion trap set to "rapid" scan rate with an AGC target of 30,000, a maximum injection time of 35 ms, and tandem MS2 data acquired starting at 136 m/z (to exclude TMTpro fragment ions). RTLS library search settings included a precursor search tolerance of 20 ppm, maximum search time of 150 ms, isotope error correction of -1/0/1/2/3. Peak selection and threshold settings included selecting *Use as a Trigger Only* and *Add Passing Search Results to Dynamic Exclusion*. The scoring threshold for acceptable library matches was a cosine score of at least 40.

Method C (FTMS Spectral Library Data Collection Method): All of the settings in this method are the same as Method A with the exception of Orbitrap MS2 acquisitions having a maximum injection time of 246 ms instead of 500 ms, and FAIMS was operated with external compensation voltage for -45 and -65 V to generate independent gas-phase fractions for each raw file.

Method D (ITMS Spectral Library Data Collection Method): All the settings in this method are the same as Method A, apart from all MS2 acquisitions being performed in the ion trap instead of the Orbitrap and FAIMS was operated with external compensation voltages for -45 and -65 V to generate independent gas-phase fractions for each raw file. The ion trap fragmentation parameters were the same as those described in Method B.

**Database Searching**

All raw files were searched using FragPipe (version 23.0) including modules MSFragger (4.3), MSBooster (1.3.9), Percolator (3.7.1), Philosopher (5.1.1), IonQuant (1.11.11), and TMT-Integrator (6.1.1). For database searching, the UniProt *Homo sapiens* protein database containing common contaminants was used (40,960 forward and decoy entries, FASTA dated 08/16/24). In the case of RTLS datasets containing high-resolution Orbitrap acquisitions and low-resolution ion trap scans (Method B), only high-resolution Orbitrap acquisitions were retained for searching within FragPipe by converting ThermoFisher raw files to mzml using MSConvert (version 3.0.25282-ce08bb4) set to filter for analyzer type “orbi”. For consistency and ensuring equitable comparisons, datasets collected with standard MS2 approach (Method A) were also converted to mzml using the same settings. For all high-resolution database searches (Methods A, B, and C), precursor mass tolerance was set to +/- 20 ppm, fragment mass tolerance was set to 20 ppm (300 ppm for low-resolution ion trap Method D), isotope error was -1/0/1/2/3, strict trypsin (cleavage at K or R) enzyme specificity was set to semi-tryptic with maximum missed cleavages of 2, peptide length of 7 to 50 amino acids, clear m/z range was set to 125.5 to 135.5 m/z, and minimum matched fragment ions was set to 4. Lysine and peptide N-terminus was set as to have a static modification of TMTpro (304.20715 Da). Variable modifications included oxidation (15.9949 Da) of Methionine residues with maximum of 3 modifications per peptide. Rescoring with deep learning prediction was performed through MSBooster with prediction of retention time and spectra. PSM validation was achieved with Percolator and protein inference was performed with ProteinProphet. For TMTpro quantification with TMT-Integrator, intensity extraction was performed with IonQuant, label type was set to “TMT-35”, non-deuterated bridge sample was set to 126, deuterated bridge reference sample was set to 135ND, mass tolerance was set to 10 ppm, minimum PSM probability was set to 0.8, minimum best peptide probability was set to 0.8, minimum purity was set to 0.5, minimum intensity was set to 5%, aggregation method was set to “median”, and ratio to abundance conversion utilized MS1. Protein-level FDR was set to 1%.

**Spectral Library generation for RTLS**

To subset the top 150 most intense fragment ions from fractionated bridge sample MS2 spectra, raw files (collected with MS Methods C and D) were converted to mzml files using MSConvert with the following command line parameters: *--filter “peakPicking vendor msLevel=1-”* , *--filter “mzWindow [136.0,2000.0]”,* and *–filter “threshold count 150 most-intense [2,2]”*. For each FAIMS fraction (-45 or -65 V) and MS2 mass analyzer

(Orbitrap or ion trap), mzml files and the “interact pep.xml” files from the FragPipe results for fractionated bridge samples were imported into SpectraST through the Trans Proteomic Pipeline (version 7.2.0) to generate an “.sptxt” spectral library using default settings. These “. sptxt” files were then converted to mzVault “.db” files using DBKey deployed in Docker Desktop. Settings for this conversion step included setting mass analyzer to “IT” and setting filter to ON for top 300 peaks (Note that the mzml files already only have the top 150 most intense ions included. Setting this filter to be above that threshold avoids an indexing issue when writing the .db file). Finally, MZvault was then used to merge the original FT and IT bridge sample “.db” files into a single “.db” for each FAIMS CV (-45 and -65 V).

**Data Importing, Normalization, and Batch Correction Steps**

Median normalized protein abundances output from FragPipe were imported into R studio (build 496) running the R programming language (version 4.5.0). Where necessary, batch effects (metadata in **Table S5**) were corrected for using the “ComBat.NA” function from the MSnSet.utils R package (version 0.2.0). For the large PBMC single-cell dataset, specific batch effects corrected for included LC column (from dual column system), N2 nanoPOTS chip, isobaric effects from carrier/bridge channel, and TMTpro channel. For dimensional reduction and algorithms requiring complete observations across all features, imputation was performed using KNN imputation (DreamAI R package, version 0.1.0) with k = 10 neighbors. To generate a reference scRNAseq PBMC atlas, 20,000 unstimulated PBMCs were randomly sampled from the dataset published by Oelen et al [5]. Next, the 20,000 cells’ scRNAseq data were processed using Seurat (version 5.3.0) by following previously described reference mapping procedures which included data normalization, harmonization, and annotation of cell types as described by Oelen et al.

**Seurat Multimodal Reference Atlas Construction**

Human PBMC scRNA-seq profiles were obtained from the 1M Immune Cell dataset (https://github.com/molgenis/1M-cells), [5] including both 10x Genomics v2 and v3 chemistries. Only unstimulated (UT) cells were retained based on metadata-derived experimental annotations. Cell type labels from the original study ("cell_type_lowerres") were used solely for later visualization and validation. Raw 10x matrices for v2 and v3 chemistries were processed separately. Seurat objects were created using cells with ≥200 detected genes. To correct for chemistry-specific mitochondrial content differences reported in the original study, we applied these QC thresholds:

- v2: cells with >8% mitochondrial reads were removed
- v3: cells with >15% mitochondrial reads were removed

In both chemistries, cells containing >9 UMIs mapping to HBB (indicative of red blood cell contamination) were excluded. This matched the filtering criteria described in the original dataset (low gene complexity, excessive mitochondrial content, or RBC contamination). For each chemistry (v2 and v3), we intersected the dataset with unstimulated barcodes and randomly sampled 10,000 cells when available. If fewer than 10,000 UT cells remained post-QC, all available cells were used. Down-sampling ensured balanced representation of chemistries and enabled computationally efficient downstream integration. Each chemistry-specific subset was independently normalized using SCTransform, and variable features were identified automatically within the

SCTransform model. PCA was run, and dimensionality inspection via elbow and loading plots informed the selection of the first 18 PCs for graph-based analysis. Cell type labels from the 1M dataset were added back by barcode intersection for visualization and to confirm expected PBMC composition. To generate a unified PBMC reference atlas, v2 and v3 SCTransformed datasets were integrated using Seurat v5. The resulting integrated object constituted the PBMC scRNAseq reference atlas used as the RNA-based reference for multimodal label transfer into the scProteomics dataset.

**Seurat Multimodal Annotation Transfer**

To enable cross-modal annotation, we first identified overlapping highly variable features between the proteomics dataset and the PBMC reference atlas by finding the overlap for the top N variable for each dataset (N = 600 for scProteomics and N = 2000 for PBMC reference atlas). Seurat's FindTransferAnchors (CCA reduction, 18 PCs) was used to establish correspondence using only these overlapping features. Transfer-based cell type predictions and per-cell confidence scores (prediction.score.max) were computed using TransferData. The number of dimensions and principal components for the reference dataset was determined using the Elbow method as implemented by the PCAtools R package (version 2.20.0). The number of dimensions assigned scRNAseq atlas-derived labels and associated annotation prediction scores were reserved for downstream steps and visualizations.

**Proteomics-driven dimensionality reduction and clustering**

To better capture proteomics-specific biological variation (particularly in proteins not represented in the scRNAseq reference atlas), downstream dimensional reduction and clustering of the scProteomic datasets was performed entirely on the top 600 variable proteins. The first fourteen PCs from this PCA were selected based on the iterative parameter testing. Neighbor graphs, Louvain clustering (resolution = 0.5), and UMAP embeddings (Euclidean metric, 13 nearest neighbors,) were constructed using these proteomics-derived PCs. This yielded six major proteomic clusters representing the structure present in the proteomics data. For each cluster, we considered only “high confidence” cells with prediction.score.max ≥ 0.56 from the Seurat annotation transfer. For every cluster × predicted cell type combination, we tested whether the number of high-confidence cells of a given predicted identity (k) exceeded expectation under the hypergeometric distribution, where:

- **N** = total number of high-confidence cells across all clusters
- **K** = number of high-confidence cells predicted as a given cell type across the dataset
- **n** = number of high-confidence cells within the cluster
- **k** = number of those cells predicted as the given cell type

One-sided enrichment p-values were computed in R as:

$$p = P(X \geq k) = \text{phyper}(k - 1, K, N - K, n, \text{lower.tail = FALSE})$$

P values were corrected for multiple hypotheses testing using the Bonferroni procedure. Only significantly enriched predictions (adjusted p-value ≤ 0.01) were accepted and clusters without a statistically significant cell type were labeled “Unknown”. Final cluster identities were then propagated to all cells within the given cluster, regardless of their individual prediction scores.

**Differential Abundance and Gene Enrichment Analysis**

$Log_2$-transformed, non-imputed protein abundance data was imported into Seurat along with the cell types identified as described above. To identify cell-type enriched proteins, each cell type was compared against all other cells using a two-sided limma/Empirical Bayes (limma R package, version 3.64.1) statistical test with *p*-values adjusted for multiple hypothesis tests using the Bonferroni correction procedure. The five lowest adjusted p-value proteins with adjusted p.value less than 0.01 and selected known marker genes were then visualized in a heatmap. Gene enrichment analysis was performed with gprofiler2 (version 0.2.3). scRNAseq data used in **Figure S9** was acquired from the Single Cell Portal.[56]

Table S1. **Median-normalized, FragPipe output for all single PBMCs**

This supplementary .csv file contains the FragPipe TMT-Integrator output that is median-normalized and rolled up to protein-level abundances. The first 10 columns describe protein metadata, after which the remaining columns provide quantification information for the TMTpro reporter ion channels in each batch analyzed. Bridge reference intensities are included with the prefix “RefInt” or “RefDInt”.

Table S2. **Single PBMC median-normalized, batch-corrected, and imputed matrix**

This supplementary .csv file provides the imputed data used for dimensional reduction and other techniques that require complete data. The first column provides the common

Gene names while every column after is for a TMT reporter ion channel and dataset corresponding to a single PBMC cell.

Table S3. **Differential abundance results across assigned cell types**

This supplementary .csv file contains four columns and describes the differential abundance results with a log2FC >= 0.5 and adjusted p.value <= 0.001. The first column (Gene) lists gene names, the second column (Cell_Type) provides the assigned cell type, the third column (log2FC) lists the log2FC, and the final column (p_value_adj) contains the adjusted p values.

Table S4. **Table listing the datasets and MS methods performed in this study.**

This supplementary .xlsx file contains three columns. The first column (Dataset) lists the names of all the raw files collected in this study, the second column, (Description) provides a short name of the experiment being performed, and the third column (MS_Method) provides the mass spectrometry method identification using the nomenclature described in **Mass Spectrometry Methodology.**

Table S5. **Analytical Metadata for all single PBMCs**

This supplementary .xlsx file contains seventeen columns. The first column represents the unique identifier for each single-cell (SampleID), followed by metadata such as Organism, Organ, Labeling_Type, Mass_Spectrometer, Raw_File_Name, and Acquisition Date. Other columns describe the N2 nanoPOTS chip used (nanoPOTS_Chip), the well of the nanoPOTS chip used (Well), the corresponding LC column of the dual column system (LC_Column), the corresponding TMTpro channel for each cell (TMTpro_32_Channel), and the TMT plex identifier (TMT_Plex_ID). Additional columns include estimated pg of peptide for each cell (Estimated_pg_peptide), the assigned cell type identified from the first pass through our informatic approach (Cell_Type_First_Pass), and the assigned cell type identified from the second pass through our informatic approach (Cell_Type_Second_Pass).

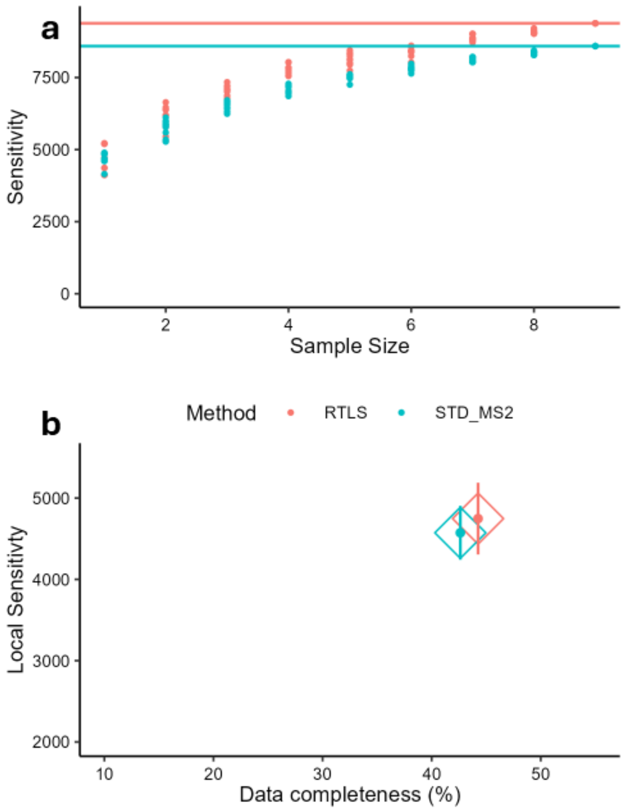

**Figure S1. Comparison of RTLS and standard MS2 method using 10 ng input.** (**a**) Cumulative sensitivity curves showing peptides identified as datasets (n = 10 total for both methods) are sampled. Solid line indicates total sensitivity (total unique peptides) (**b**) Mean local sensitivity (average number of peptides per dataset) compared to overall data completeness. Lines represent +/- 1 standard deviation.

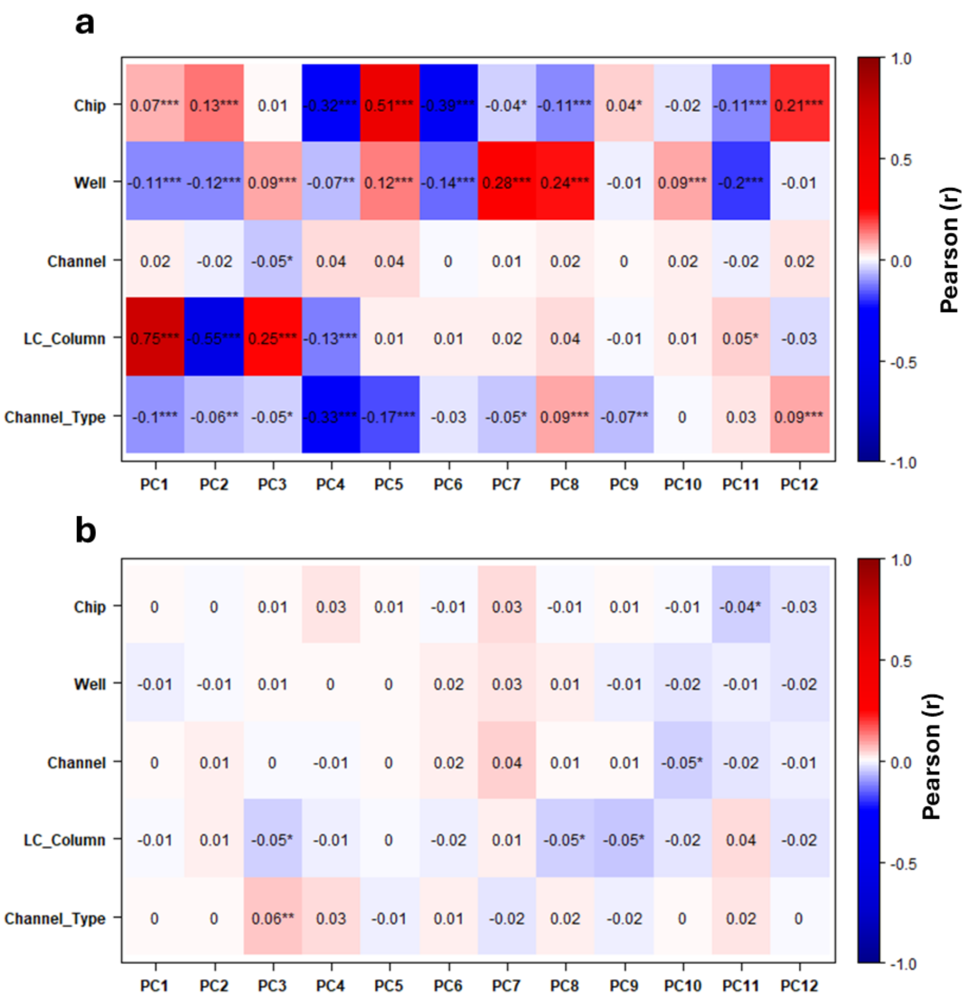

**Figure S2. Eigencor plots showing the correlation of batch effects with PCs.** (**a**) Pearson correlations (r) between the first twelve principal components (x-axis) and several variables known to have batch effects (y-axis), before batch correction and (**b**) after batch correction. “Chip” refers to the physical N2 nanoPOTS platform used, “Well” refers to the well location, “Channel” refers to the TMTpro channel, “LC_Column” refers to the LC column on the dual column system (1 or 2), and “Channel_Type” refers to TMTpro channels that are either isotopically adjacent to the higher input bridge channels or non-adjacent.

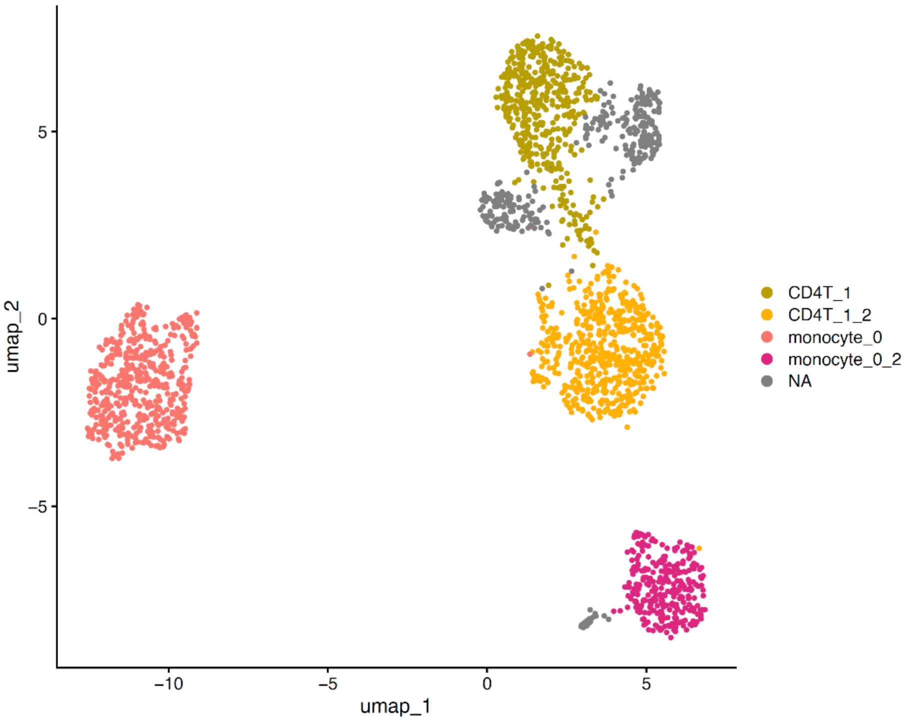

**Figure S3. UMAP of identified cell types from initial cell typing analysis.** Colors indicate different cell types identified using the informatic pipeline described in methods. UMAP embedding was produced using the first 11 PCs from the scProteomics data, nearest neighbors = 27, and min.dist set to 0.1. Underscore followed by a number indicates the underlying cluster from which the cell type was mapped to.

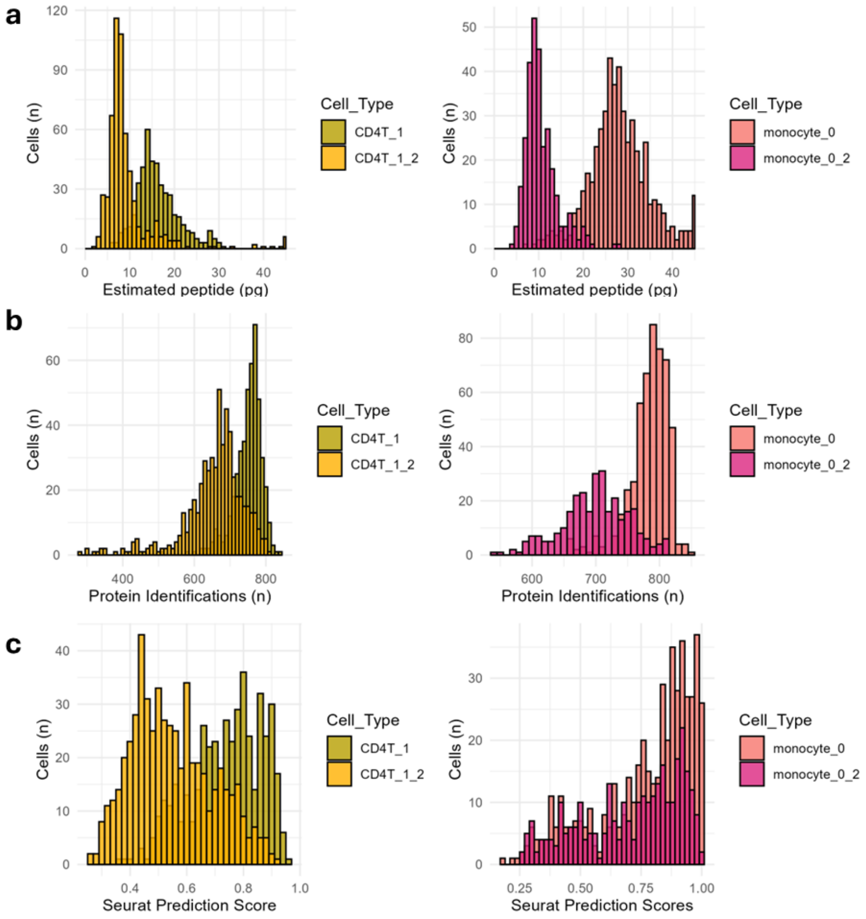

**Figure S4. Histograms comparing estimated peptide input, identified proteins, and Seurat prediction scores for CD4+ T-cells and monocyte clusters. (a)**. Estimated peptide input for each single cell. This was calculated by dividing the median cell reporter ion intensity by its respective median bridge channel intensity, followed by multiplying the estimated bridge input amount (150 pg). (**b**) Total protein identifications for each single cell. (**c**) Maximum prediction scores provided by Seurat for each single cell.

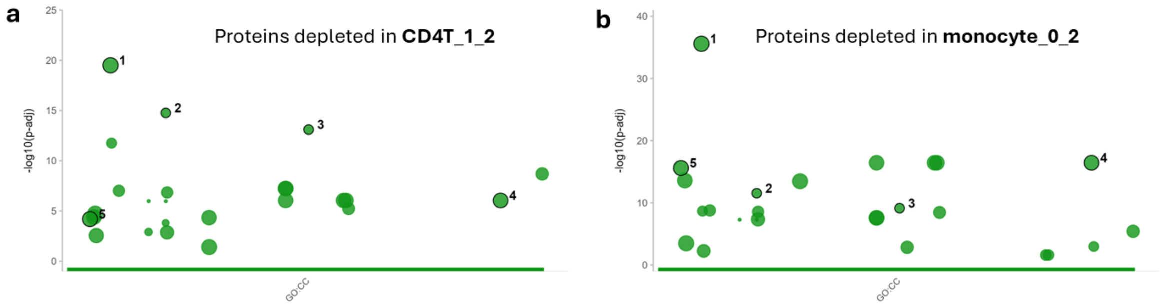

| id | source | term_id | term_name | term_size | p_value |
|---|---|---|---|---|---|
| 1 | GO:CC | GO:0005829 | cytosol | 893 | 3.1e-20 |
| 2 | GO:CC | GO:0022626 | cytosolic ribosome | 80 | 1.7e-15 |
| 3 | GO:CC | GO:0044391 | ribosomal subunit | 82 | 7.9e-14 |
| 4 | GO:CC | GO:1903561 | extracellular vesicle | 665 | 9.0e-07 |
| 5 | GO:CC | GO:0005576 | extracellular region | 764 | 6.4e-05 |

g:Profiler (biit.cs.ut.ee/gprofiler)

| id | source | term_id | term_name | term_size | p_value |
|---|---|---|---|---|---|
| 1 | GO:CC | GO:0005829 | cytosol | 893 | 2.6e-36 |
| 2 | GO:CC | GO:0022626 | cytosolic ribosome | 80 | 2.8e-12 |
| 3 | GO:CC | GO:0044391 | ribosomal subunit | 82 | 7.2e-10 |
| 4 | GO:CC | GO:1903561 | extracellular vesicle | 665 | 3.6e-17 |
| 5 | GO:CC | GO:0005576 | extracellular region | 764 | 2.5e-16 |

g:Profiler (biit.cs.ut.ee/gprofiler)

**Figure S5. Gene enrichment analysis of CD4+ T-cell and monocyte clusters.**

Manhattan plot of gene ontological cellular compartment terms highlighting shared terms between cell types. (**a**) shows GO terms found for CD4T cells while (**b**) shows GO terms found for monocytes. Differential abundance analysis was performed with limma prior to gene enrichment of features with statistical significance (adjusted p.value < 0.01) and negative log2FC.

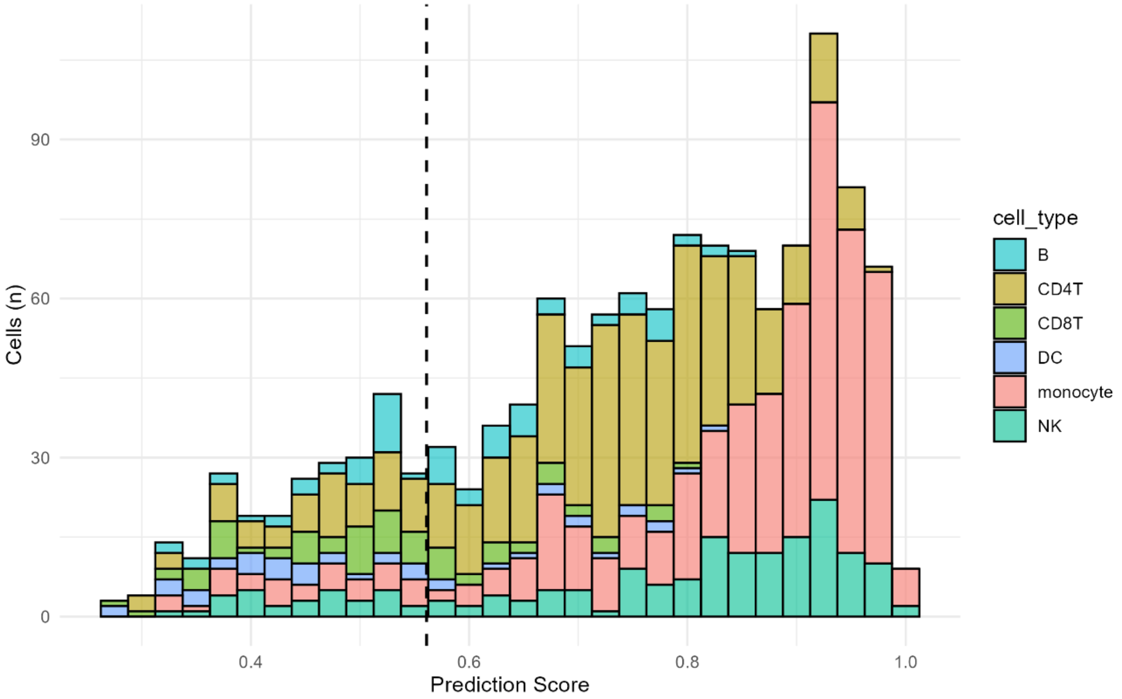

**Figure S6. Histogram of Seurat prediction scores for 1,275 PBMCs.** Histogram of the Seurat prediction scores for the indicated cell types. Dashed vertical line indicates the prediction score threshold of 0.56 for considering an annotation as “high confidence”. “DC” = dendritic cell, “NK” = natural killer cell, and “B” = B-cell

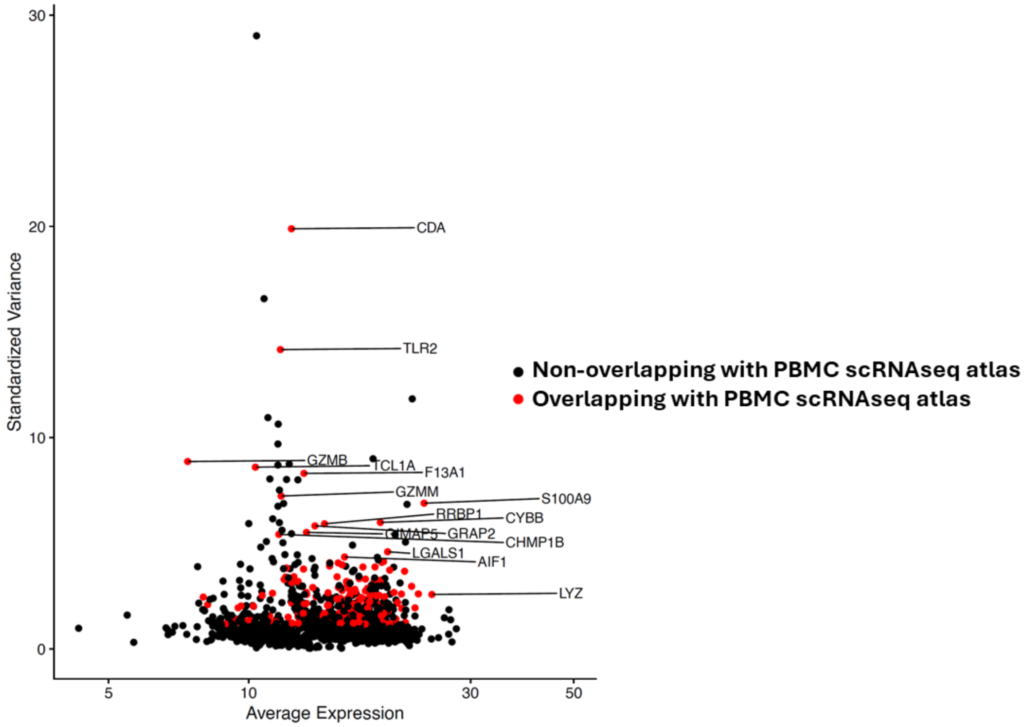

**Figure S7. Variable feature plot of 1,648 proteins measured in 1,275 PBMCs.** X-axis shows the average log2(Intensity) for each protein while y-axis presents the standardized variance determined using variance stabilizing transformation (VST). Points colored in red indicate overlapping variable features between scProteomic and scRNAseq data.

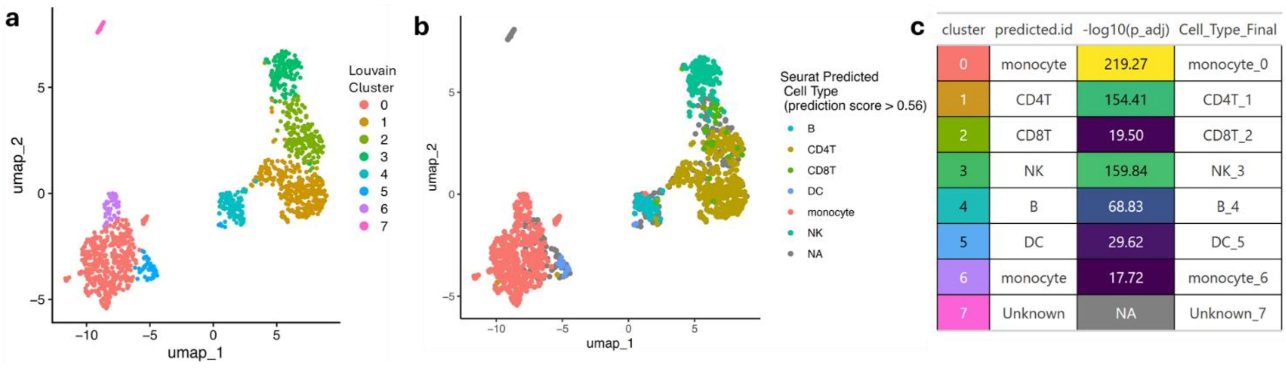

| cluster | predicted.id | -log10(p_adj) | Cell_Type_Final |
|---|---|---|---|
| 0 | monocyte | 219.27 | monocyte_0 |
| 1 | CD4T | 154.41 | CD4T_1 |
| 2 | CD8T | 19.50 | CD8T_2 |
| 3 | NK | 159.84 | NK_3 |
| 4 | B | 68.83 | B_4 |
| 5 | DC | 29.62 | DC_5 |
| 6 | monocyte | 17.72 | monocyte_6 |
| 7 | Unknown | NA | Unknown_7 |

**Figure S8. Louvain clustering, Seurat predicted, and final annotated cell types.** (**a**) UMAP highlighting the Louvain clusters identified using the shared nearest neighbor graph constructed from the PCA of the 350 highest varying proteins from the scProteomic data. (**b**) Same UMAP as (**a**), but with colors indicating the high-confidence cell type annotations predicted by Seurat (prediction score > 0.56) (**c**) Table describing the Louvain cluster, Seurat predicted cell type for that cluster, hypergeometric test adjusted p-value associated with the cluster x cell type, and the final annotation naming ("Cell_Type_Final"). "DC" = dendritic cell, "NK" = natural killer cell, "B" = B-cell, and "NA" = not applicable.

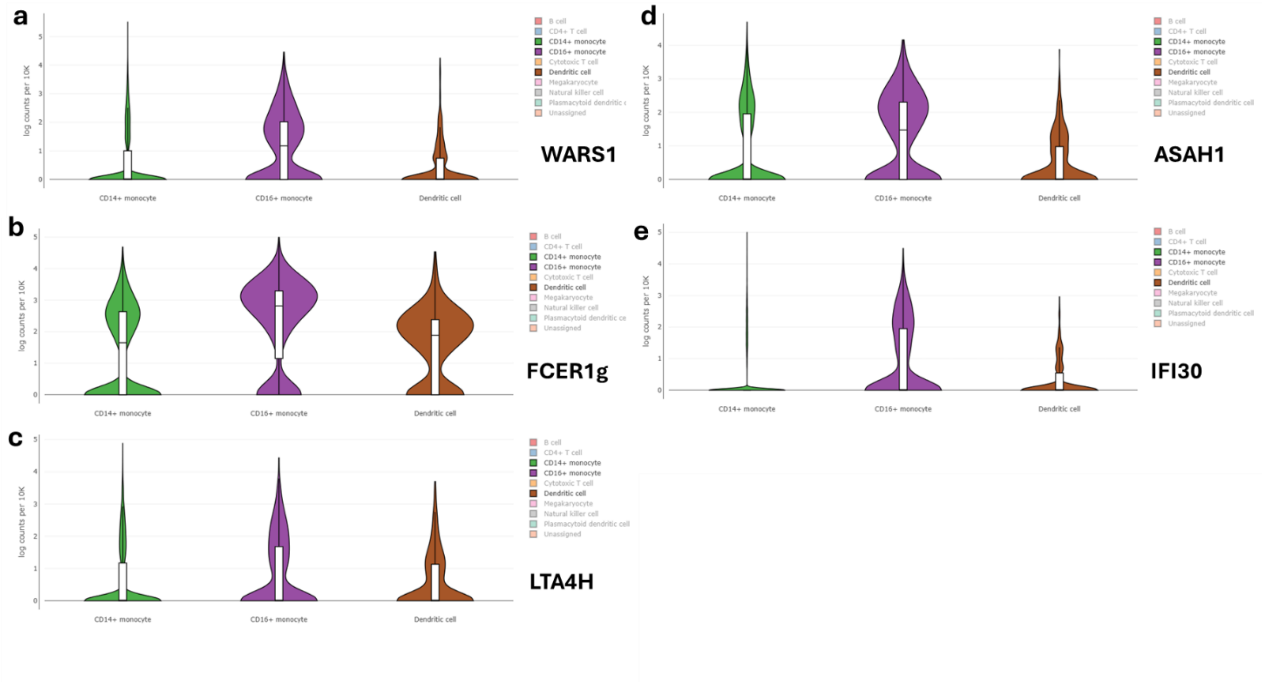

**Figure S9. scRNAseq expression data for CD16+ monocyte enriched proteins.** Violin plots of scRNAseq expression data for monocyte lineage cells. Each plot corresponds to proteins found enriched for the monocyte_6 cluster in the scProteomic data and includes (**a**) WARS1, (**b**) FCER1G, (**c**) LTA4H, (**d**) ASAH1, and (**e**) IFI30. Colors indicate CD14+ monocytes (green), CD16+ monocytes (purple), and dendritic cells (brown).
